# Hive-associated bacteria improve honeybee survival during *Paenibacillus larvae* infection

**DOI:** 10.64898/2025.12.24.696342

**Authors:** Anuja Shrestha, Yva Eline, Kenzie Sundstrom, James T. Van Leuven

## Abstract

Honeybees (*Apis mellifera* L.) are the most common pollinators of crops. Honeybee larvae are susceptible to *Paenibacillus larvae*, a spore-forming Gram-positive bacterium that causes American Foulbrood (AFB), one of the most destructive brood diseases worldwide. Existing antibiotic treatments are undesirable due to increasing pathogen resistance and their residual accumulation in bee products. Consequently, there is increasing interest in biological agents and natural strategies for AFB control. However, most studies remain limited to *in vitro* or *in vivo* experiments and rarely evaluate impacts on adult bees. In this study, we systematically investigated hive-associated bacteria for their potential to enhance larval survival during *P. larvae* infection. Bacteria isolated from AFB-infected combs were sequenced and screened for antagonistic activity. Several *Bacillus* species—including *B. zhangzhouensis*, *B. subtilis*, *B. amyloliquefaciens*, *B. licheniformis*, and *B. mojavensis*—inhibited one or more *P. larvae* strains. Notably, the lysate of *B. licheniformis* suppressed all tested strains; Further characterization revealed that its main antibacterial component consists of heat-stable proteins between 30–50 kDa. Two *Bacillus* strains were selected for larval infection assays using laboratory-reared honeybee larvae: both significantly improved survival by 42% and 71%, respectively. To evaluate potential effects on adult bees, newly emerged workers were caged and fed *B. licheniformis* ASx lysate. Their survival differed from that of untreated controls but remained comparable to the Terra-Pro-fed bees, a Terramycin-based AFB treatment, and their gut microbiome remained similar to that of untreated controls. Overall, our findings suggest that hive-associated *Bacillus* species offer promising, low-impact candidates for AFB disease management.

## INTRODUCTION

Managed honeybees (*Apis mellifera* L.), one of the most important crop pollinator species, provide around 50% of global crop pollination (Kleijn et al., 2015) and play a significant role in global food production, contributing to approximately one-third of the total human food supply (Khalifa et al., 2021). In the United States, managed honeybees alone add $15 billion worth of crop pollination services each year (Kulhanek et al., 2017; Morse & Calderone, 2000). However, beekeepers have consistently experienced high losses of managed honeybee colonies each year due to a plethora of biotic and abiotic risk factors (Hristov et al., 2020). One of the causes of honeybee illness and colony collapse is American foulbrood disease (AFB).

AFB is an important bacterial disease in honeybee larvae that has spread globally and is recognized as an economically important disease for apiculture. It is caused by *Paenibacillus larvae* (*P. larvae*) (Firmicutes), a spore-forming, gram-positive, highly specialized pathogen with only one established host, the honeybee larvae (Forsgren et al., 2010; Nowotnik et al., 2024).

The bacterium infects young larvae via the ingestion of contaminated food, after which the bacterial spores germinate, and the vegetative bacteria propagate rapidly and proliferate in the midgut before they breach the epithelium and invade the hemocoel (Ghorbani-Nezami et al., 2015; Yue et al., 2008), causing cell injury and larval death. The dead larvae degrade to a ropy thread when drawn out with a matchstick, which, when dried, creates scales containing millions of spores readily transmissible between colonies and across hives by foraging bees or apiculturists (Genersch 2010). These spores are highly heat-resistant and stable for up to 35 years, making the control of AFB very difficult (Genersch 2010).

AFB has been particularly devastating because of inadequate current treatment options. Burning colonies and contaminated hive materials are expensive for beekeepers, resulting in the loss of both hive materials and productive hives. Alternatively, the use of antibiotics, such as oxytetracycline or tylosin, to clear the infection is commonly practiced. However, they are not sufficient to combat the disease due to increased pathogen resistance to the treatments, and the accumulation of residuals in honeybee products raises concerns (Al-Ghamdi et al., 2018; Oliveira et al., 2015; Tian et al., 2012). These antibiotics are not permitted for use in European countries, as current legislation prohibits the inclusion of pharmacotherapy treatment, and the only control options are the physical elimination of infected colonies (Nowotnik et al., 2024).

Additionally, it has been reported that these antibiotics can inhibit members of the bee gut microbiota and cause individual bees to starve (Baffoni et al., 2021; Raymann et al., 2017). This has led to an increasing concern as well as interest in the investigations of alternative, effective, and novel control methods of AFB, including several biological control agents with various modes of action, which could be used for apiculture.

Different studies from different parts of the world have reported that certain bacterial species isolated from apiarian sources showed antagonistic activity against *P. larvae* and could be an alternative and effective control of AFB disease in honeybees (Alippi & Reynaldi, 2006; Minnaard & Alippi, 2016; Sabaté et al., 2009; Yoshiyama & Kimura, 2009). A few other studies reported some bacterial strains not related to any apiarian sources: strains from GenBank previously shown to have probiotic potential (Nowotnik et al., 2024), a strain from the USDA GenBank (Cherif et al., 2008), a strain isolated from the native soil from the Brazilian Atlantic Forest (Benitez et al., 2012), were reported to inhibit *P. larvae*, suggesting their potential for use in the control of AFB disease in honeybees. However, all these studies were only conducted in the lab using *in vitro* assays. A study from Saudi Arabia investigated the effects of gut bacterium, including *Lactobacillus kunkeii* and *B. licheniformis,* on the mortality percentage of honeybee larvae infected with *P. larvae*, where they reported that the addition of these bacteria to the normal diets was able to significantly reduce the mortality percentage, suggesting the antagonistic potentialities against AFB disease in honeybees (Al-Ghamdi et al., 2018). However, a systematic investigation of hive-associated bacteria against *P. larvae* in *in vitro*, *in vivo*, and in adult bees has not been explored yet.

Considering that *in vitro* results can or cannot be replicated *in vivo*, we aim to comprehensively investigate the potential of hive-associated bacteria in improving the survival of honeybee larvae during *Paenibacillus larvae* infection. First, we isolated bacteria from AFB-infected honeybee hive samples and taxonomically identified them. Secondly, we tested their ability to inhibit *P. larvae* strains *in vitro*. Third, the potential positive candidates from the *in vitro* assays were further examined for their inhibitory ability using laboratory-reared honeybee larvae. Finally, the best bacterial candidate was selected to test its ability to impact the survival of adult bees as well as to investigate its impact on the adult bee gut microbiome, using caged adult worker honeybees in the laboratory.

## MATERIALS AND METHODS

### 1. Source and isolation of bacterial strains

*Paenibacillus larvae* strains Y-3650 (NRRL B-3650) and 368 (ATCC-25368) were acquired from the University of Nevada, Las Vegas (Yost et al., 2016), and 25747 (ATCC-25747) was received from the American Type Culture Collection (ATCC). Among these strains, Y-3650 and 25747 belong to Enterobacterial Repetitive Intergenic Consensus (ERIC) group I, whereas 368 belongs to ERIC group IV (Paull et al., 2024). The remaining *P. larvae* strain [ONT-R-1C (strain 55)] and all the *Bacillus* strains tested for inhibition assays were isolated in our lab, either extracted from bee guts from our apiary in Moscow, Idaho, or from the AFB-infected hives samples from Pullman, Washington, USA, in 2023. The *Lactobacillus* strains used in this study were received from NCIMB GenBank (NCIMB 15254) (Figure 3).

For culturing, the acquired *P. larvae* and *Lactobacillus* strains were streaked from their respective glycerol stocks from −80°C freezer onto 1mg/L of thiamine hydrochloride Brain Heart Infusion (BHI) agar (Catalog#DF0418-17-7, BD DifcoTM, USA) plates and de Man, Rogosa and Sharpe (MRS) (Catalog#1.10660, GranuCult prime, EMD Millipore Corporation) media plates with 5% fructose, respectively. The remaining strains were isolated in our lab. Under sterile conditions, both the adult bee guts and infected beehive samples were ground in mortars and pestles in 1mL of sterile Phosphate buffer at pH 7 (PBS, Catalog#BP399-1, Fisher Scientific, USA). The ground mixture of adult bee guts and infected beehive samples was then serially diluted (10^0^ to 10^-4^) with PBS, and 100 µL of each dilution was plated onto MRS and BHI media, respectively. The BHI plates were incubated at 37°C with 5% CO2 for 72 hours, and the MRS plates were incubated in an anaerobic chamber under the same temperature. A single unique colony from the plates was streaked onto new BHI or MRS plates, and incubated again for 3 more days, followed by making cultures by taking a single clean colony from the streaked plates in 3 mL of BHI or MRS broths, and incubated at 37°C in a shaking incubator at 120 rpm for 24 to 48 hours depending on the turbidity of the cultures. Once the cultures were ready, 1 mL was used to prepare a pair of glycerol stocks with 40% glycerol and stored at −80 °C, and half was used for DNA extraction.

### 2. Identification of bacterial strains

One to two days old bacterial liquid cultures, depending on their turbidity (1mL each), were used for genomic DNA extraction using the Quick-DNA HMW MagBead Kit (Catalog #D6060, Zymo Research), following the manufacturer’s protocol. The DNA samples were quantified using a Qubit^TM^ Flex Fluorometer (Catalog#Q33327, Invitrogen Life Technologies, CA, USA), followed by 16S rRNA amplification. The PCR Master Mix working stock was prepared by mixing 5 µL 5X reaction buffer, 0.5 µL 12.5 mM dNTPs, 0.250 µL Q5 High-Fi DNA polymerase, 17 µL molecular grade water, 1.25 µL of 10 µM 27f primer (5’-AGAGTTTGATCMTGGCTCAG-3’), and 1.25 µL of 10 µM 907r primer (5’-CCGTCAATTCMTTTRAGTT-3’). A total final volume of 25 µL per reaction was prepared by adding 24 µL of the Master mix and 1 µL of bacterial genomic DNA. The PCR amplification conditions were as follows: initial denaturation at 98°C for 1 min, followed by 30 amplification cycles of (98°C for 30 sec, 58°C for 30 sec, 72°C for 1 min), and a final extension at 72°C for 5 min, using a T100^TM^ Thermal Cycler (Serial# 621BR76793, Bio-Rad Laboratories, Inc. Singapore). The amplified PCR products were verified by gel electrophoresis and purified using the GeneJET PCR Purification kit (Catalog#FERK0701, ThermoFisher Scientific, USA). The DNA was then re-quantified using Qubit and sent for 16S rRNA sequencing at Eurofins Genomics, DNA Sequencing Lab, Louisville, USA. The 16S DNA sequences were trimmed at the 5’ and 3’ ends and aligned to construct contigs, and BLAST searched to obtain taxonomic predictions, using Geneious Prime Software® 2025.1.2 (https://www.geneious.com). The highest percent identity matches with the higher query coverage and were counted as the top matches.

For the *P. larvae* strain [ONT-R-1C (strain 55)], DNA was sent to Plasmidsaurus for whole-genome sequencing. The sequences of all these strains were submitted to GenBank, and their accession numbers can be found in Bioproject PRJNA1377638.

### 3. *In vitro* assay for screening the inhibitory activity against *Paenibacillus larvae* strains

For the assay, *P. larvae* strains were streaked from glycerol stock onto 1mg/L of thiamine hydrochloride Brain Heart Infusion (BHI) agar (Catalog#DF0418-17-7, BD Difco^TM^, USA) plates and incubated at 37 °C for 72 hours. A single colony from the plates was used to make a liquid culture and incubated at 37 °C for 24 hours at 200 rpm. 100 µL of each culture was added to 3 mL of BHI top agar, mixed, and then poured onto a BHI agar plate to make a bacterial lawn of *P. larvae* strains. The *Bacillus* and *Lactobacillus* strains were streaked from their respective glycerol stocks onto BHI and MRS plates, respectively, and incubated as mentioned above. A single colony from the respective plates was used to make bacterial cultures and incubated for 24 hours (*Bacillus* strains) and 48 hours (*Lactobacillus* strains) in a 37°C shaker incubator at 120 rpm. Both bacterial cultures, as well as lysates, were used to make spots on the lawns of *P. larvae* strains. For this, half of the bacterial cultures were centrifuged at 100 rpm for 2 minutes, and the resulting supernatants were filtered through 0.22 µM membrane filters to obtain lysate from each bacterial strain. Two different methods were used for screening these bacterial strains for inhibitory activity *in vitro* against *P. larvae* strains. For testing the bacterial cultures, wells were punched on the *P. larvae* lawns, and 100 µL of the cultures was inoculated into the wells.

For testing the lysates, 20 µL was directly spotted on the bacterial lawns. All the plates were incubated at 37 °C for 24 hours with the lids up. The bacterial inhibition by bacterial cultures or lysates was determined by checking the halos of inhibition zones formed on a bacterial lawn on a solid agar plate. Among these strains, only *B. licheniformis* (ONT-R2-ASx) was selected for further *in vitro* studies.

### 4. Testing different sizes of membrane filters to collect lysates from *Bacillus licheniformis* - ONT-R2-ASx to determine different lysate sizes that have antimicrobial effects

Liquid culture of ONT-R2-ASx was prepared as described above (see Method section 3). Then, a subculture was prepared using the overnight culture and BHI broth at a 1:10 ratio in a sterile flask and incubated for 24 hours. The bacterial mixture was then centrifuged at 1000 rpm for 3 minutes, after which the lysate was filtered first through a 0.45 μM membrane filter, followed by using a 0.22 μM membrane to obtain the lysate. Some of the 0.22 μM lysate was then passed through a 100 kDa molecular-weight cut-off (MWCO) membrane filter and likewise, through 50 kDa and 30 kDa MWCO filters, respectively (Figure 1). Then, a total of four different lysate sizes: (A) Less than 0.22 µM-greater than 100 kDa; (B) Less than 100 kDa - greater than 50 kDa; (C) Less than 50 kDa -greater than 30 kDa; (D) Less than 30 kDa and their 1:10 dilutions, were tested against the *P. larvae* strains Y-3650, 368, and strain 55 by spotting 20 µL onto their bacterial lawns. Each lysate size and its one-fold (1:10) dilutions were screened independently in triplicate. The extent of bacterial inhibition was determined by measuring the diameter of the inhibition zone formed on the bacterial lawn on a solid agar plate.

**Figure 1.**
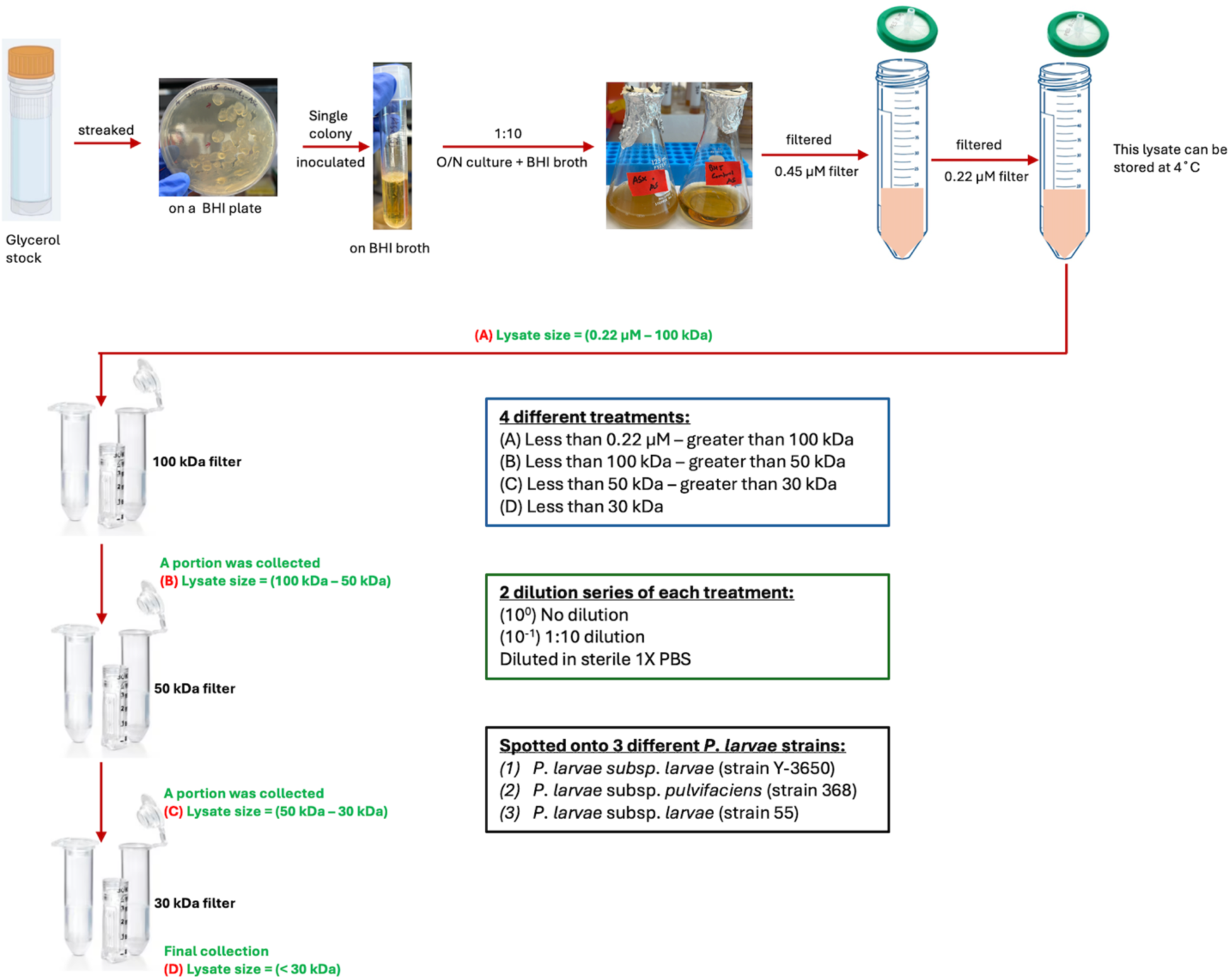
Flow diagram showing the steps from streaking the ONT-R2-ASx strain from its glycerol stock and collecting different sizes of lysates from its cultures, which were then spotted onto the lawns of three *P. larvae* strains.

### 5. Testing the effect of proteinase K, royal jelly, and temperature on the antimicrobial activity of *Bacillus licheniformis* - ONT-R2-ASx (50-30 kDa) lysate

The ONT-R2-ASx (50-30 kDa) lysate was further used for stability tests to understand its peptide nature. The susceptibility to proteolytic enzyme (proteinase K), royal jelly (Feed A, fed to the larvae during laboratory rearing), and heat treatment was evaluated. To investigate the effect of proteinase K on the antimicrobial activity of the lysate, it was treated with proteinase K at a concentration of 100 μg/mL at 37°C for 30 mins, after which the treated lysate was tested for its antimicrobial activity against *P. larvae* strains (Kim et al., 2020; Epparti et al., 2022). Since the lysate was planned to be fed to honeybee larvae with royal jelly (Feed A) on the day of grafting, it was important to determine whether Feed A could affect the lysate’s antimicrobial activity. Therefore, the lysate and Feed A (1:1) were incubated at 37°C for 30 mins, and this mixture was spotted on the lawns of the *P. larvae* strains. To analyze thermal stability, the lysate was serially three-fold diluted (10^0^ to 10^-3^), and each dilution was heat-treated at 100°C for 10 mins. The heat-treated tubes were kept in ice immediately before spotting them onto the bacterial lawns.

Then, 20 μL of each of the above-mentioned treatments was spotted on the lawns of *P. larvae* strains as mentioned above. After incubation for 24 hours, the diameter of the inhibition zone formed on the bacterial lawn was measured. Each treatment was screened independently in triplicate, then analyzed using One-Way ANOVA, and means were compared using Tukey’s pairwise mean comparison within each *P. larvae* strain, using SAS software (The SAS 9.4M9 and SAS Studio v3.83) with a significance level set at p=0.05.

### 6. Infection assays using laboratory-reared honeybee larvae

We did six different larval infection combinations using laboratory-reared honeybee larvae. (1) using either Y-3650 or 368 *P. larvae* strains or their combination; (2) 2D cultures with *P. larvae* strain Y-3650, (3) NCIMB *Lactobacillus* strain’s culture with Y-3650, and (4) *B. licheniformis*- ONT-R2-ASx (50-30 kDa) lysate, hereafter called ‘ASx lysate’, with the *P. larvae* strain we cultured in our lab (Strain 55).

Honeybees’ colonies were maintained at the Soil Steward Farm, Research Station at the University of Idaho, USA. Healthy, aged-matched, first instar larvae from hives established in 2024 and 2025 were used for the experiments. The frames with larvae were covered with damp paper towels and were transported to the laboratory in a foam nuc-box with hot water bottles to maintain temperature and humidity. Larvae were grafted using Chinese grafting tools, and each larva was reared in an individual queen cup placed into a 48-well cell culture plate. The grafting tools, as well as the cups with culture plates, were UV-sterilized before use. Before the larval grafting, the queen cups were supplemented with a mixture of 10 µL pre-warmed artificial larval diet “Feed A” (44.25% royal jelly, 5.3% glucose, 5.3% fructose, 0.9% yeast extract, and 44.25% water) and treatment-bacterial inoculum. Each cell culture plate included a negative control group (n=12) consisting of uninfected larvae fed with control feed A and three other treatments. The larvae were fed the following treatments on the day of grafting (Day 0): controls at 1:10 with BHI broth: Feed A; pathogens treatments at bacterial cells target 100 CFU/µL: Feed A; antibacterial strain treatments at 1:10 bacterial cells target 100 CFU/µL.: Feed A; and mixtures of pathogen and antibacterial strain treatments at cells target 100 CFU/mL: Feed A. For the ONT-R2-ASx (50-30) kDa lysate experiment, the same ratio; however, non-diluted lysate was used. The larvae were fed equal amounts of Feed A on day-0 and day-1 whereas the feed amount (42.95% royal jelly, 6.4% glucose, 6.4% fructose, 1.3% yeast extract and 42.95% water) was gradually increased from days 2 to 5 with 20 µL on day 2, 30 µL on day 3, 40 µL on day 4, and 50 µL on day 5 as described by Crailsheim et al (2012). The temperature and humidity of the larva-rearing incubator were maintained at 35°C and 95-100%, respectively, throughout the rearing period. The plates were monitored and photographed daily; dead larvae were recorded and removed. The difference in the survival of the tested larvae throughout the experiments was analyzed by using log-rank tests with all pairwise comparisons with Benjamini-Hochberg multiple test correction. All the statistical analyses were performed and compared in R using the “survival” and “survminer” packages. The value P ≤ 0.05 was considered the threshold for significance.

### 7. Testing the effect of *B. licheniformis* ONT-R2-ASx (50-30 kDa) lysate in adult worker honeybees reared in laboratory conditions

The larval experiment results showed that the ASx lysate could improve the survival of honeybee larvae during AFB infection. However, the lysate’s effect on adult bees’ health is not known. It is essential to have scientific evidence to support their efficacy because the results from the larval experiment may or may not be fully replicated in adult bees. Therefore, to determine the effectiveness of the lysate on adult bees and to ensure the survival of honeybee colonies, we performed cage experiments under laboratory conditions with newly emerged adult bees.

For this study, sixteen healthy brood frames containing capped brood were collected from our research apiary located at the University of Idaho (Moscow, Idaho, USA) and transferred to the laboratory in nucleus colony (Nuc) boxes. Any larvae or honey was scraped off using sterile scrapers, and the frames were placed in the laboratory incubator at 33°C and 60% relative humidity for 24 hours until adult bee emergence. On the experimental day 0 (D0), the newly emerged adult (NEWs) worker bees were caged in plastic insect cages sized 20 x 14.5 x 14.5 cm: 100 bees per cage. They were maintained at 33°C and 60% RH, and different treatments were provided on the same day, D0. Comb materials were provided in all cages to mimic the field environment. In total, 13 cages were used in the experiment: 4 treatments with 3 replicate cages per treatment, and one cage was used for measuring evaporation rates; everything else was kept the same, except that it was without bees. Among the cages with treatments, they were fed with 10 mL control (40% sugar syrup only) (n=3); 10 mL 40% sugar syrup mixed with laboratory-grade tetracycline (Catalog T252525G, TCI America ^TM^, Fisher Scientific, USA) at the doses of 0.1 mg/mL (n=3); 10 mL 40% sugar syrup mixed with commercial Terra-Pro mediated Terramycin treatment (MLL: 25-055; Item#DC-550; Mann Lake Ltd. Hackensack MN) at the doses of 0.014 gram/mL; and ONT-R2-ASx (50-30 kDa) lysate with 40% sugar syrup at the ration of 1:10. These treatments were administered three times at the interval of 5 days at D0, D6, and D12 (Figure 2). All cages received ad libitum control 40% sugar syrup between treatments, and their consumption was recorded at 24-hour intervals throughout the experiment. To explore the impact of the treatments on adult worker bees, both visual and microbiome assessments were conducted. In terms of visual evaluation, the survival of bees in the cages was recorded daily (D1 to D17). The survival differences among bees across treatments were analyzed using log-rank tests, with all pairwise comparisons corrected for multiple testing using the Benjamini-Hochberg method. All statistical analyses were performed and compared in R using the “survival” and “survminer” packages, with P ≤ 0.05 as the threshold for significance.

**Figure 2.**
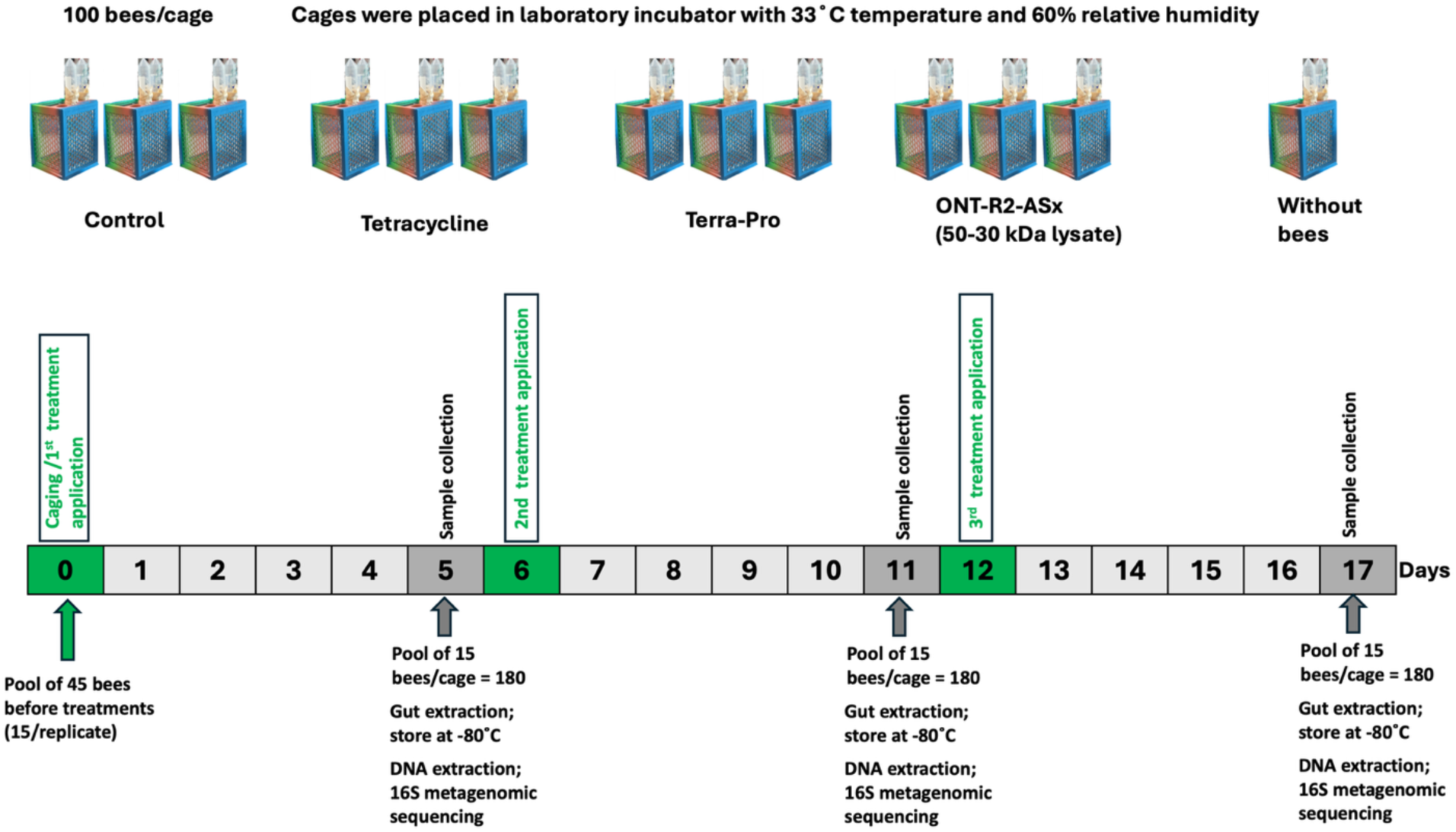
Study design and methodology used for testing the effect of ONT-R2-ASx (50-30 kDa) lysate, including lab-grade tetracycline and commercially available Terra-Pro in newly emerged worker honeybees in cages under lab conditions. The study was conducted in September-October 2025.

**Figure 3.**
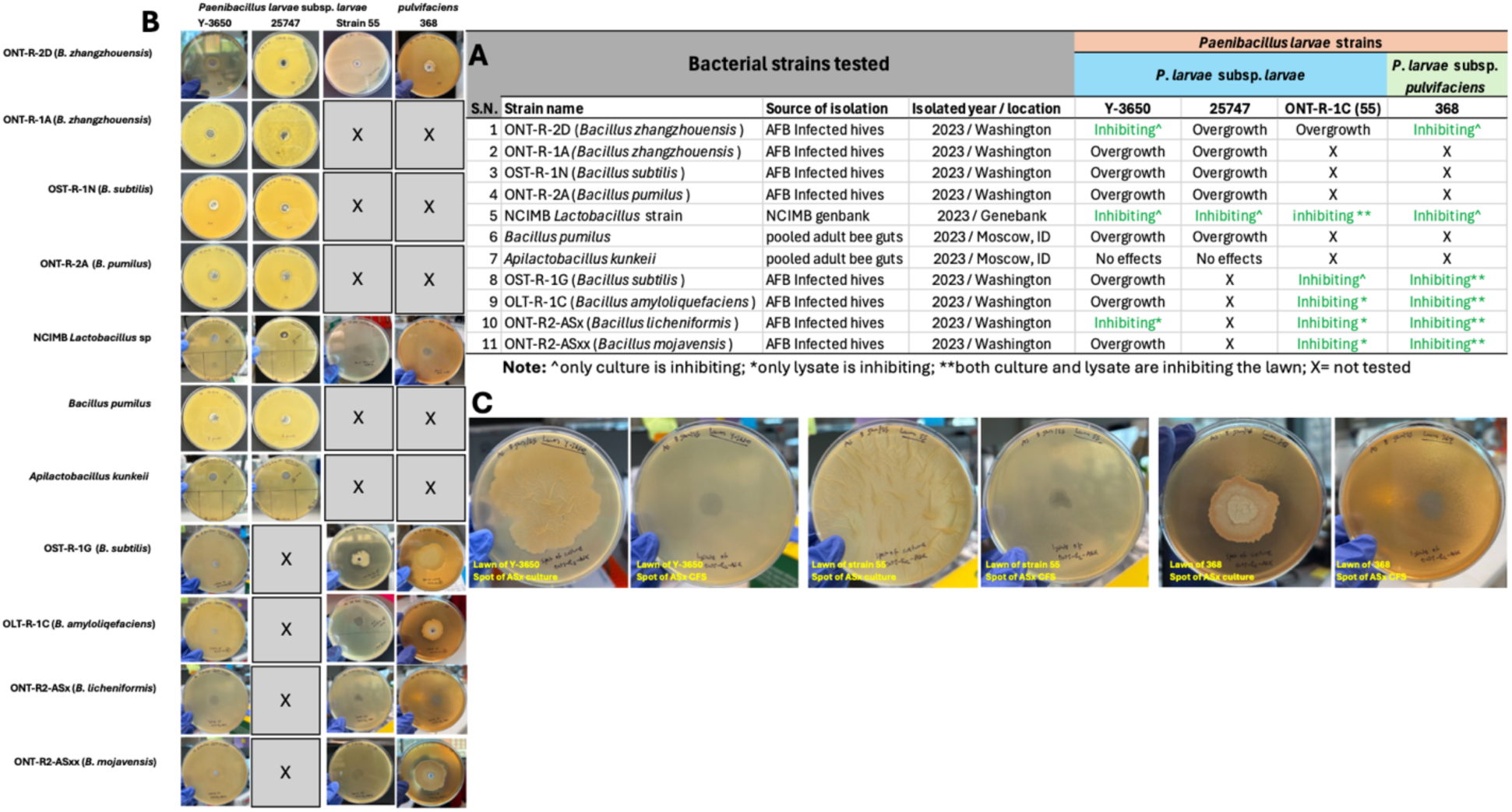
Bacterial strains showing the potential to inhibit *Paenibacillus larvae* strains *in vitro*: (**A**) a list of bacterial strains used in the study, their sources of isolation, isolated year, locations from where the samples were received, and their effects on *P. larvae* strains. (**B**) Representative plate pictures showing the effects of the tested strains against the *P. larvae* strains. (**C**) Example showing the effect of cultures and lysates of the strain *B. licheniformis* (ONT-R2-ASx) against the *P. larvae* strains.

The gut microbiome assessment study was conducted to evaluate the effects of different treatments on gut microbiota community composition, how these effects change across treatments, how the treatments affect gut microbiomes, and how similar or different their effects are. Additionally, it would be interesting to understand and compare the gut microbiomes of sterile NEWs that were used in the experiments, before any treatment application, for baseline microbiome information. Since the comb materials were provided to all cages to mimic the field environment, it was necessary to identify the microbial composition of the added comb samples and determine if they had any impacts on the microbiome composition of the treatments.

Additionally, it would be useful to evaluate the impact of laboratory cage rearing on the bee gut microbiome. For this, we included a mark-recapture treatment group. First, three pools of 15 NEWs per pool were removed on Day 0 (D0) before the treatment application for the baseline microbiome data. Secondly, on the same day (D0), around 200 NEWs were marked and returned to the hives, which were later collected from the hives (n=3 pools of 6 bees) to extract guts.

Thirdly, pools of 15 bees per cage were removed on days 5, 11, and 17. Finally, on day 17 (D17), the comb materials from all cages were collected and made 3 pools together (n=3), with a total of 44 samples. After each pool of bees, guts were extracted and stored at −80°C until DNA extraction was performed. The DNA was extracted using the ZymoBIOMICS DNA Microprep kit (Catalog # D4301, Zymo Research) according to the kit’s protocol.

The samples were sequenced by Plasmidsaurus using Oxford Nanopore (ONT) R10.4.1 flow cells with long-read sequencing libraries constructed using v14 library prep chemistry and in-house 16S gene primers. Taxonomic identification was completed by Plasmidsaurus using emu (v3.5.1) against rrnDB (v5.6) and NCBI targeted Loci databases (https://www.plasmidsaurus.com/). All the sequences were submitted to GenBank and can be found in Bioproject PRJNA1377638.

All statistical analyses were conducted in R (v4.5.2) using custom scripts. Taxonomic data were imported and analyzed using phyloseq (v1.54.0). Beta diversity was assessed via principal coordinates analysis (PCoA) with Bray-Curtis dissimilarity matrices, and community similarity was calculated using Jaccard distances with the vegan package (v2.7.2). Differential abundance of treatments, ASx lysate, Terra-Pro, and tetracycline was performed using two complementary methods: DESeq2 (v1.50.2) and ANCOM-BC2 (v2.12.0), with control as the reference group (Lin & Peddada, 2024). Differential abundance testing of hive-reared and controls was also performed with DESeq2 and ANCOM-BC2, with comb as the reference group. Significance thresholds were set at adjusted p-value (padj) <0.05 for DESeq2 and q-value <0.05 for ANCOM-BC2.

## RESULTS

### 1. Bacterial strains isolated from the bee guts and the AFB-infected hives, with their potential to inhibit *Paenibacillus larvae in vitro*

We isolated and characterized, by 16S rRNA sequencing, many strains of bacteria from honeybee guts and hive debris. These included *Bacillus zhangzhouensis*, *B. subtilis*, *B. pumilus*, *B. amyloliquefaciens*, *B. licheniformis*, *B. mojavensis, Apilactobacillus kunkeei,* and a *Paenibacillus larvae* strain [ONT-R-1C (strain 55)] (Figure 3A). These *Bacillus* and *Lactobacillus* strains were tested for their ability to inhibit four different *P. larvae* strains: Y-3650, 25747, ONT-R-1C (strain 55), and 368.

We found that some strains of the bacteria that we isolated can strongly inhibit the *P. larvae* when co-cultured on agar plates, some have overgrowth on the lawns, and some do not have any effect at all (Figure 3A, B). Overgrowth was characterized by the expansion of the “inhibitory” bacteria from the inoculation spot. This made it impossible to tell if *P. larvae* was growing as well because we could only see the growth of the *Bacillus* or *Lactobacillus* species on the plate. It is interesting to note that in some cases, one bacterial strain had different effects on different *P. larvae* strains. When we compared inhibition from cell cultures to inhibition from spent media (cell-free lysates), we found that it was most common that *P. larvae* was inhibited by both the culture and the cell-free lysate. The second most common occurrence was for *P. larvae* to be inhibited by only a culture of the bacteria we tested or for only the lysate to inhibit *P. larvae* growth. For example, the culture of *Bacillus zhangzhouensis* (Strain ONT-R-2D) can inhibit Y-3650 and 368; however, it had overgrowth over 25747 and ONT-R-1C (strain 55) lawns. Both *B. pumilus* isolates that we tested had overgrowth on *P. larvae* lawns. NCIMB *Lactobacillus* strain and *B. licheniformis* had inhibitory effects on the tested *P. larvae* strains, either with their cultures (cells), with lysates (without cells), or with both cultures and lysates. Among all the tested bacterial strains, *B. licheniformis* (ONT-R2-ASx) seemed to have interesting inhibitory effects. Only the lysate showed the zones of inhibition on the lawns of Y-3650 and strain 55, whereas the cultures had overgrowth on those lawns. However, both its lysate and culture showed inhibitions on the lawn of 368 (Figure 3C). Therefore, we selected the lysate of strain ONT-R2-ASx for further *in vitro* study to further characterize the components causing the inhibitory activities.

### 2. The lysate size 50-30 kDa from the strain *Bacillus licheniformis* - ONT-R2-ASx has the antimicrobial factor that inhibits *P. larvae* strains, the causative agent of AFB disease in honeybees

Four different sizes of ONT-R2-ASx lysates were spotted onto the lawns of *P. larvae* strains Y-3650, 368, and strain 55 (Figure 1). Both the non-diluted and 1:10 dilutions of each lysate were tested against the *P. larvae* strains. The spot plate assays showed that all the lysate sizes (non-diluted) except those less than 30 kDa showed inhibition against the *P. larvae* lawns (Figure 1, Figure 4, and Table 1). In terms of diluted lysates (1:10), only two different lysate sizes (0.22 μM-100 kDa) and (100-50 kDa) showed inhibitions, however, only on the lawn of strain 55 (Table 1, Figure 4). The antimicrobial effect was consistent across all lysate sizes except for less than 30 kDa lysate (Table 1), indicating that the primary antimicrobial compound should have a molecular weight under 50 kDa and above 30kDa. The ONT-R2-ASx lysate was able to pass through a 30 kDa molecular weight cut-off (MWCO); however, it didn’t show inhibition zones on the *P. larvae* lawns. We noticed a reduction in the size of the zone of inhibition as filter size decreased, indicating that perhaps more than one component of the lysate is responsible for inhibition, with some components being stopped as the filters become progressively more restrictive.

**Figure 4.**
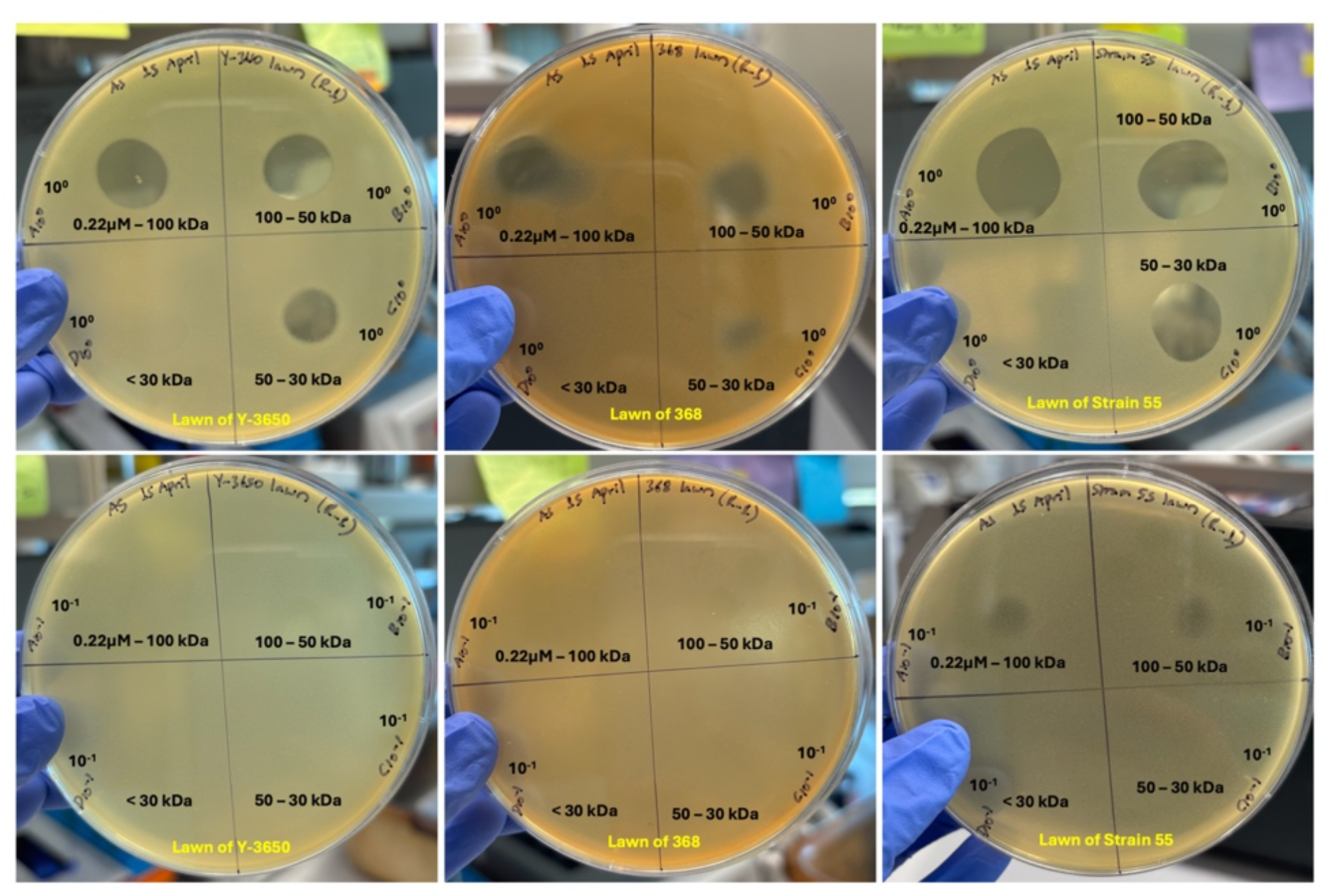
Testing different ONT-R2-ASx lysates on the lawn of three different *Paenibacillus larvae* strains. Lysate sizes (A) Less than 0.22µM-greater than 100 kDa; (B) Less than 100 kDa - greater than 50 kDa; (C) Less than 50 kDa -greater than 30 kDa; (D) Less than 30 kDa. Two dilutions of each lysate size were spotted: top row - (10^0^) Not diluted lysate; and bottom row - (10^-1^) 1:10 diluted lysate

**Table 1.** Testing the ability of different sizes of ONT-R2-ASx lysates against different strains of *Paenibacillus larvae*. Diameter (Mean ± SEM) of inhibitory zones obtained in the study. Data were calculated from three independent experiments.

| *P. larvae* strains | Dilution of lysate | Inhibition zone diameter (in cm) (Mean ± SEM) | | | |
| --- | --- | --- | --- | --- | --- |
| | | 0.22μM-100kDa | 100-50kDa | 50-30kDa | Less than 30kDa |
| Y-3650 | 10<sup>0</sup> | 1.6 ± 0 | 1.4 ± 0 | 1.1 ± 0 | 0 |
|  | 10<sup>-1</sup> | 0 | 0 | 0 | 0 |
| 368 | 10<sup>0</sup> | 1.5 ± 0.1 | 1.15 ± 0.15 | 0.6 ± 0 | 0 |
|  | 10<sup>-1</sup> | 0 | 0 | 0 | 0 |
| Strain 55 | 10<sup>0</sup> | 2.05 ± 0.05 | 1.85 ± 0.05 | 1.55 ± 0.05 | 0 |
| | 10<sup>-1</sup> | 0.6 ± 0 | 0.6 ± 0 | 0 | 0 |

### 3. Influence of Proteinase K enzyme, royal jelly, and temperature on the antimicrobial activity of *Bacillus licheniformis* - ONT-R2-ASx (50-30 kDa) lysate

To further investigate the components of the ONT-R2-ASx lysate causing antimicrobial activity against *P. larvae*, we exposed the lysate fractions to various treatments and measured any losses in antimicrobial activity. The antimicrobial activity was lost when the ONT-R2-ASx lysate was exposed to proteinase K at a concentration of 100 μg/mL (Table 2). This suggested that the antimicrobial activity of the lysate from ONT-R2-ASx is mediated mainly by proteins in the 30-50 kDa size range that function as antimicrobial peptides (AMPs). The antimicrobial activity of the tested lysate was not significantly affected when the lysate was treated with royal jelly, and it was also resistant to heating at a temperature of 100°C, since its zones of inhibition were not significantly different before and after the treatments, except for the treatment with proteinase K (Table 2; Figure 5; Figure S1). This showed that the AMPs associated with the ONT-R2-ASx (50-30 kDa) lysate are heat-stable and are not degraded by the royal jelly.

**Figure 5.**
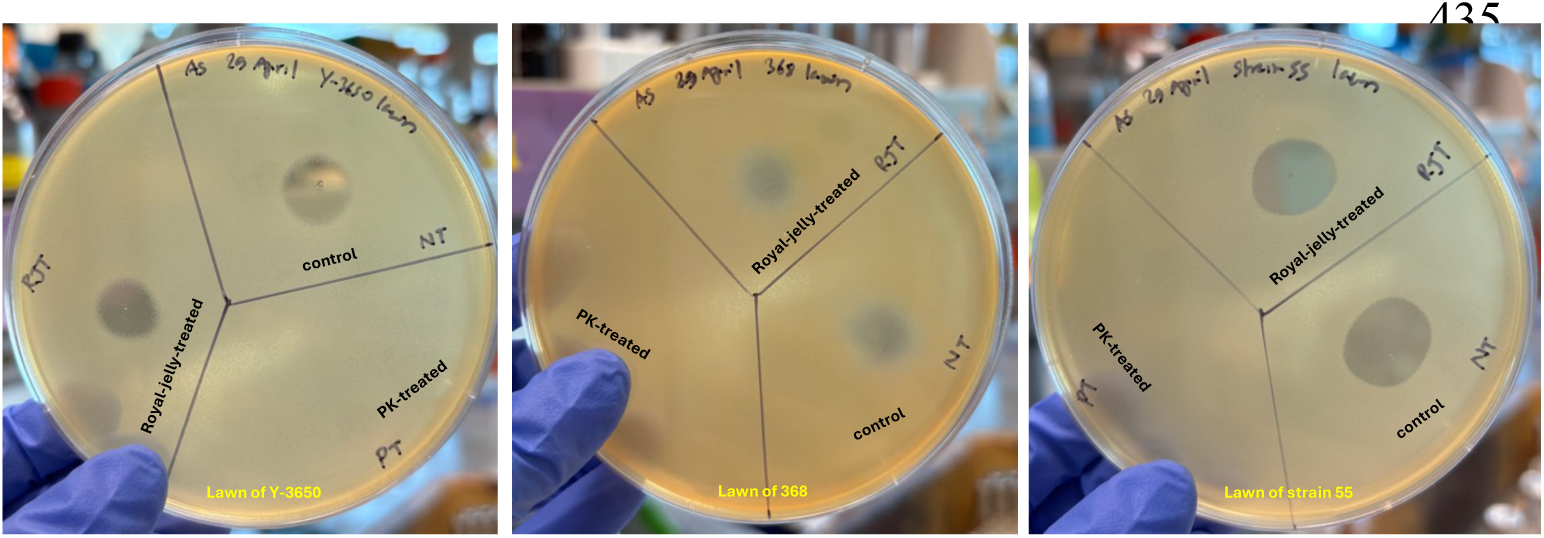
Representative plates of spot plate assay of different treatments of ONT-R2-ASx (50-30 kDa) lysate on the lawn of *P. larvae* strains. Pictures of heat-treated spot assays are shown in the supplementary file: Figure S1.

**Table 2.** Testing the ability of different treatments of *B. licheniformis* - ONT-R2-ASx (50-30 kDa) lysates to suppress *Paenibacillus larvae* strains.

| Treatment | Inhibition zone diameter (in cm) (Mean ± SEM) | | |
| --- | --- | --- | --- |
|  | Y-3650 | 368 | Strain 55 |
| ASx 50-30 kDa Control | 1.1 ± 0.00 a | 0.65 ± 0.05 a | 1.45 ± 0.05 a |
| ASx 50-30 kDa PK-treated | 0 b | 0 b | 0 b |
| ASx 50-30 kDa RJ-treated | 1.0 ± 0.00 a | 0.55 ± 0.05 a | 1.40 ± 0.10 a |
| ASx 50-30 kDa Heat-treated | 1.05 ± 0.05 a | 0.45 ± 0.05 a | 1.45 ± 0.05 a |

LS-means with the same letter are not significantly different ( $P \leq 0.05$ ). PK-treated = proteinase K-treated; RJ-treated = royal jelly-treated

**Table 3.** Jaccard distances quantifying dissimilarities in bacterial community composition between sample types.

|  | NEWs | Control | Tetracycline | Terra-Pro | ASx lysate | Hive-reared | Comb |
| --- | --- | --- | --- | --- | --- | --- | --- |
| NEWs | 0 | - | - | - | - | - | - |
| Control | 0.65 | 0 | - | - | - | - | - |
| Tetracycline | 0.667 | 0.471 | 0 | - | - | - | - |
| Terra-Pro | 0.657 | 0.382 | 0.269 | 0 | - | - | - |
| ASx lysate | 0.708 | 0.372 | 0.537 | 0.5 | 0 | - | - |
| Hive-reared | 0.654 | 0.613 | 0.429 | 0.5 | 0.692 | 0 | - |
| Comb | 0.722 | 0.5 | 0.581 | 0.531 | 0.524 | 0.542 | 0 |

**Notes:** Values range from 0 (identical communities) to 1 (different communities). Antimicrobial treatments, tetracycline and Terra-Pro, show high similarity to each other, reflecting similar effects on community composition. ASx lysate maintains greater similarity to untreated controls than to antimicrobial treatments, consistent with minimal microbiome disruption. The NEWs differ substantially from all developed microbiomes, reflecting their undeveloped state. Hive-Reared bees show moderate distances from all caged treatments, indicating environmental influences on microbiome composition beyond initial colonization. These distance metrics complement the PCoA ordination patterns and support treatment-specific effects on community structure.

### 4. The virulence levels of two different ERIC-type strains of *Paenibacillus larvae* on the laboratory-reared honeybee larvae

Two different strain types, Y-3650 (ERIC type I) and 368 (ERIC type IV), were used to infect laboratory-reared honeybee larvae to determine their virulence levels. Approximately 100 cells of each of these strains were fed to larvae and monitored for 5 days. The results showed significant differences in the survival percentages of larvae, infected with Y-3650 [P=3.0e-05, log-rank test with BH correction, n=864, df=3] and 368 [P=4.9e-06, log-rank test with BH correction, n= 864, df=3] when compared with the larvae treated with broth controls. However, there is no significant difference in the percentage larval survival when infected with Y-3650 and 368 [P=0.59, log-rank test with BH correction, n= 864, df=3] (Figure 6A-E, I.1).

**Figure 6.**
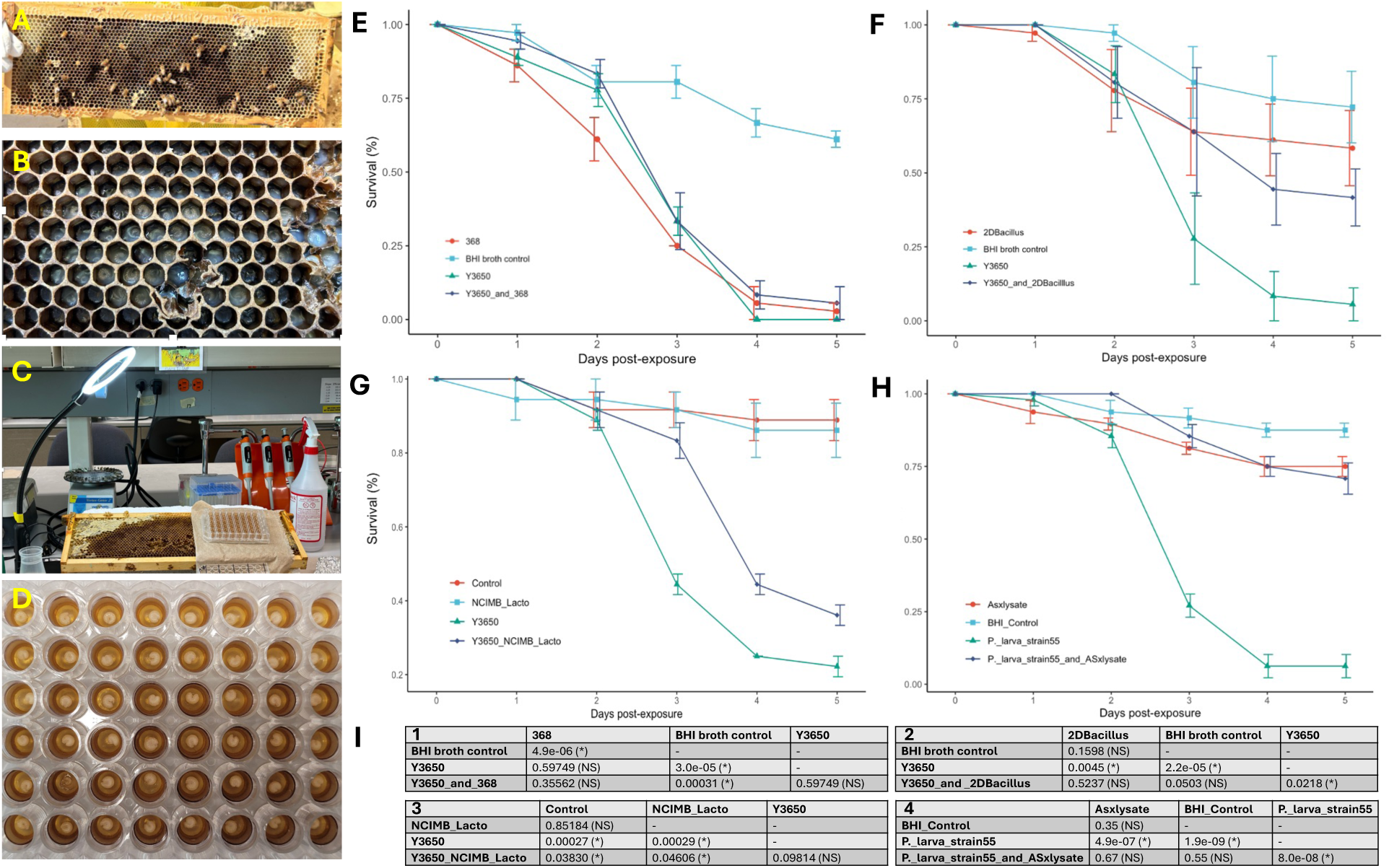
Testing the potential of different antimicrobial strains or lysate in the laboratory-reared honeybee larvae against the AFB pathogen, *P. larvae* strains. (A) A-frame collected from our apiary in Moscow, Idaho; (B) Frame with first-instar larvae; (C) Lab set-up for larval grafting; (D) Larvae in a 48-well plate after 24 hours of grafting; (E) Results of the larval experiment using two different *P. larvae* strains Y-3650 and 368; (F) *Bacillus zhangzhouensis*-2D and *P. larvae* strain Y-3650; (G) NCIMB *Lactobacillus* strain and *P. larvae* strain Y-3650; and (H) Results showing the ONT-R2-ASx lysate could improve the survival of honeybee larvae during the infection by *P. larvae* strain 55. Standard error bars are shown (I) Results of the statistical analyses of all the larval experiments above, in R using the “survival” and “survminer” packages (1) Y-3650 and 368; (2) 2D and Y-3650; (3) NCIMB *Lactobacillus* strain and Y-3650; and (4) ONT-R2-Asx lysate and *P. larvae* strain 55.

### 5. The effects of *Bacillus zhangzhouensis* 2D and NCIMB *Lactobacillus* strains’ culture on the laboratory-reared honeybee larvae

The cultures of both strains—the NCIMB *Lactobacillus* and *B. zhangzhouensis* strain 2D—showed inhibitory effects in the *in vitro* assays. Therefore, these strains were tested in honeybee larvae experiments to evaluate their effectiveness against the *P. larvae* strain Y-3650.

The larval experiments using 2D and Y-3650 showed no significant differences in larval survivals when larvae were fed only 2D cultures (58%) and both Y-3650 and 2D together (42%), when compared with the ones fed with only BHI broth (72%) [P=0.15] and [P=0.05] respectively. However, they showed significant differences in survival when fed with these bacteria when compared with the larvae inoculated only with Y-3650 (6%) [P=0.0045] and [P=0.021], respectively (Figure 6A-F, I.2). This result supported the *in vitro* results that strain 2D has the potential to inhibit *P. larvae* growth.

The larval survival results were somewhat different in the case of the NCIMB *Lactobacillus* strain and Y-3650. The *Lactobacillus* strain showed similar effects in the larval survival as the BHI controls [P=0.85] but showed a significant survival difference from those larvae fed with both *Lactobacillus* and Y-3650 [P=0.04], suggesting that the *Lactobacillus* strain alone did better than with co-inoculation with Y-3650. In addition, the survival percentage of the larvae with the co-inoculation with the *Lactobacillus* strain with Y-3650 had no significant difference from the larvae fed only with Y-3650 [P=0.09] (Figure 6-G, I.3). This contradicts the results of the NCIMB *Lactobacillus* strain having the potential to inhibit Y-3650 *in vitro*.

### 6. The antimicrobial peptide of *Bacillus licheniformis* - ONT-R2-ASx (50-30 kDa) lysate could improve the survival of honeybee larvae during *P. larvae* infection

As reported earlier, ONT-R2-ASx (50-30 kDa) has an antimicrobial compound that can inhibit *P. larvae* strains in *in vitro* assays. To determine if these antimicrobial effects are relevant to honeybees, infection assays were conducted on laboratory-reared honeybee larvae. The larval experiment results showed that most [88% (42/48)] of the media control-treated larvae survived. When compared with the control-treated larvae, we found no significance difference in survival of larvae infected with *P. larvae* strain 55 and treated with the ONT-R2-ASx (50-30 kDa) lysate [70.8% (34/48)] [P=0.55] or when only treated with the lysate [75% (36/48)] [P=0.35] (Figure 6A-H, I.4). The larvae infected with *P. larvae* strain 55 only, had poor survival, with only 6.25% (3/48) remaining at the end of the experiment. This was significantly different when compared to larvae treated with ONT-R2-ASx (50-30 kDa) lysate [P= 4.9e-07, log-rank test with BH correction, n=1152, df=3] or infected with *P. larvae* strain 55 and treated with the ONT-R2-ASx (50-30 kDa) lysate [P=8.0e-08, log-rank test with BH correction, n=1152, df=3].

### 7. The potential prophylactic use of *B. licheniformis* ONT-R2-ASx (50-30 kDa) lysate in honeybee hives and its potential impacts on the adult worker honeybees

The larval experiments showed that *B. lichceniformis* ONT-R2-ASx (50-30 kDa) - ASx lysate protected honeybee larvae during *P. larvae* infection *in vitro*. To determine the effect of the lysate on the health of adult bees, we performed cage experiments where bees were fed with either sugar syrup or syrup with either tetracycline, Terra-Pro, or the lysate.

On Day 5 post-first inoculation, the survival percentage of the adult bees treated with ASx lysate (94.4%) and Terra-Pro (95%) was significantly different from that of the non-treated negative control bees (97.4%; P=0.016, log-rank test with BH correction, n=6600, df=3), whereas the survivability percentage of tetracycline-treated bees (97.6%) was similar to the controls (P=0.192, log-rank test with BH correction, n=6600, df=3) (Figure 7A). On the same day (D5), 15 bees were pooled out from each cage for microbiome analysis, and on Day 6, the second dose of the same treatments was fed to the remaining bees. On Day 11 post-second inoculation, similar results were obtained as of Day 5; there was no significant difference in the survival percentage of adult bees treated with ASx lysate (67.6%) versus Terra-Pro (72.3%) [P=0.965, log-rank test with BHI correction, n=13200, df=3]; however, both treatments had significantly different survival percentages when compared with controls and tetracycline-treated bees (91.8% and 85.1% respectively) [P=1.1E-14, 6.6E-13 respectively] (Figure 7B). Again, 15 bees were pooled per cage on Day 11, and the third dose of the same treatments was fed to the leftover bees. On Day 17 post-third inoculation, the results were different. All three treatments had significantly different bee survival percentages than the non-treated control bees. Also, the survival percentages of ASx lysate and Terra-Pro-treated bees were significantly different than that of tetracycline-treated bees (Figure 7C).

**Figure 7.**
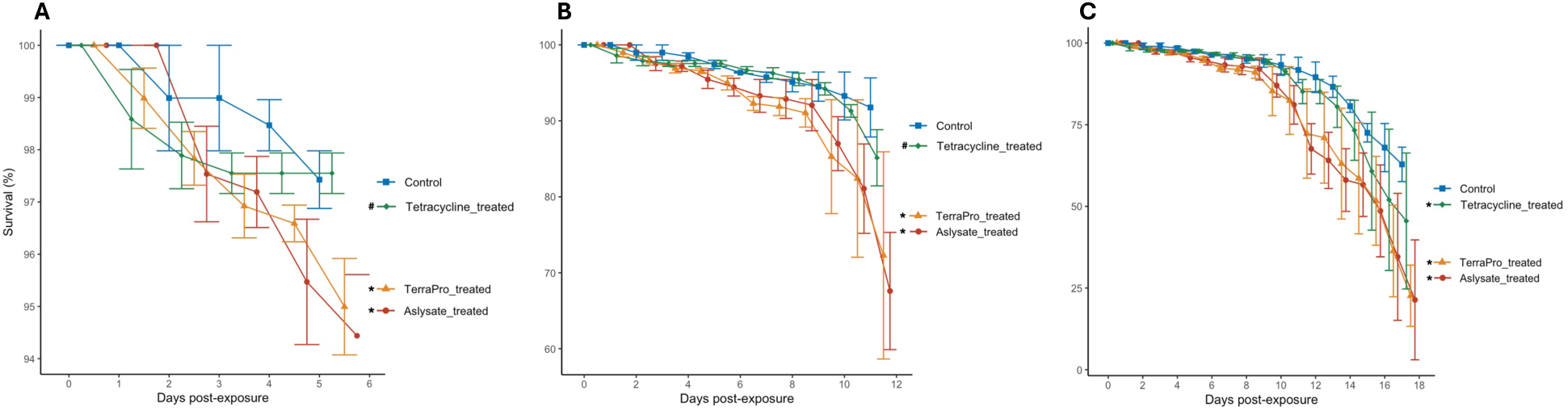
Mean percentage survival of newly emerged adult worker bees (n=300/treatment) in the laboratory cages, fed with a total of 3X treatments of either tetracycline or TerraPro, or ASx lysate. (A) Bees caged received 1X treatments, from 0 to 5 days; (B) received 2X treatments from 0 to 11 days; (C) received 3X treatments from 0 to 17 days. # indicates the survival percentage is not significantly different from the control, and * indicates the survival percentage is significantly different from the control with P <0.05.

### 8. The impact of antimicrobial treatments on the adult honeybee microbiome

We sequenced a total of 44 pools of dissected bee guts using full-length 16S rRNA sequencing. The mean number of post-quality filtering reads was 4,824 ± 625 per sample with a mean quality score of 20.79 and a mean read length of 1,444 base pairs. Taxonomic classification identified 54 unique taxa comprising ≥ 0.1% relative abundance across all samples (Supplemental file: sequencing_stats_all). The overall taxonomic composition reflected the characteristic core microbiome of adult honeybee workers, with *Lactobacillus* and *Gilliamella* as the dominant genera. *Lactobacillus* was the most abundant genus in 40 of 44 samples, represented by 7 distinct species. In the four samples where *Lactobacillus* was not the dominant genus, *Gilliamella* (n=2), *Serratia* (n=1), and *Tatumella* (n=1) were the most abundant instead.

However, *Lactobacillus* remained present as the second-ranked genus in these samples. These four samples were distributed across treatment groups and timepoints (NEWs, Day 5 Control, Day 5 Terra-Pro, and Day 17 ASx lysate) (Supplemental file: dominant_genus). *Bifidobacterium*, a core member of the honeybee gut microbiome, was not identified in any of the samples, including hive-reared and comb (Figure 8; Supplemental file: core_bacteria_presence).

**Figure 8.**
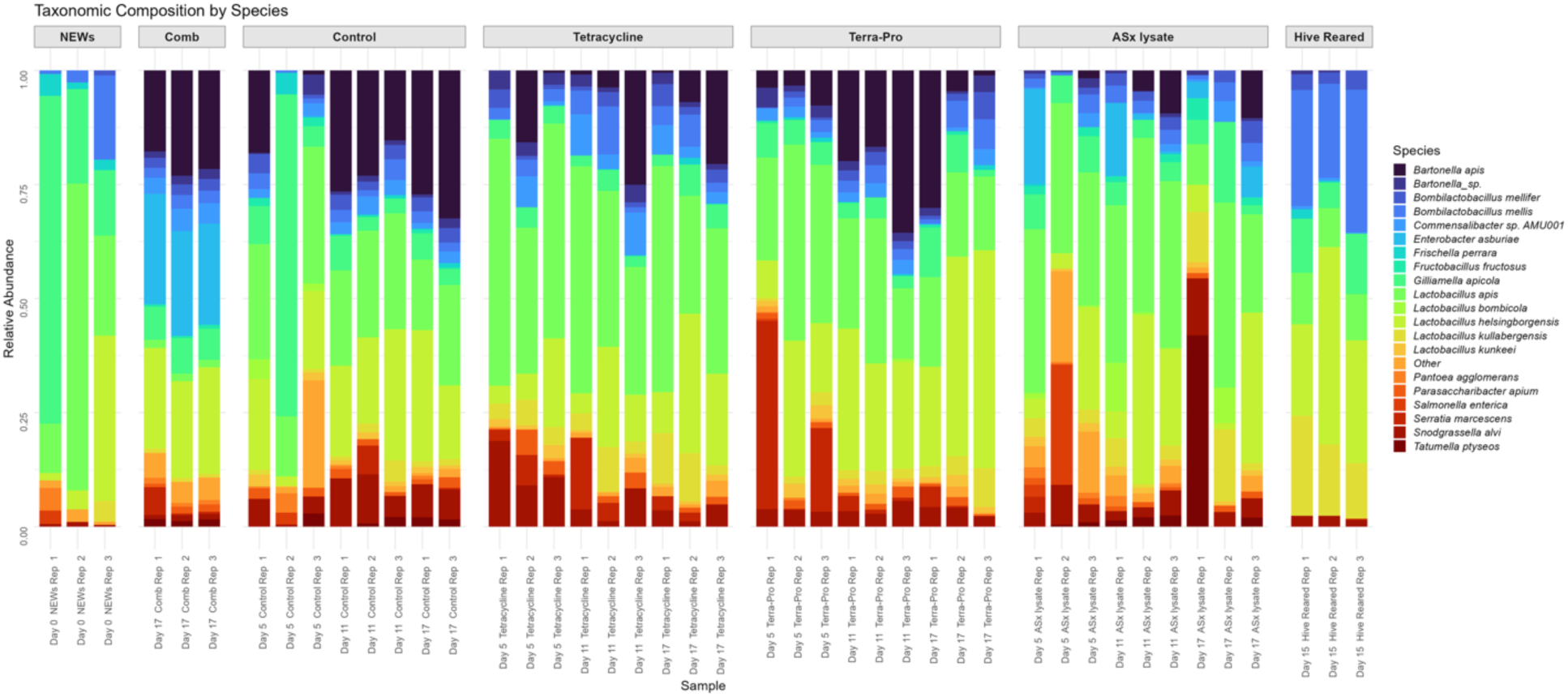
Taxonomic composition by species: Relative abundance of the top 20 most abundant bacterial species across all samples, grouped by treatment and sampling day. Each bar represents an individual sample, with colors indicating different bacterial species. Sample types are organized left to right: NEWs (Newly Emerged Workers, collected on Day 0 for baseline estimate), comb (bacterial source material, collected on Day 17 for comparison), control (caged, untreated), tetracycline (caged, antibiotic treatment), Terra-Pro (caged, antibiotic treatment), ASx lysate (caged, lysate treatment), Hive-Reared (NEWs marked on Day 0 and returned to hive, collected on Day 15 comparison). Taxonomic composition remains dominated by *Lactobacillus* and *Bombilactobacillus* species across treatments, with Gilliamella as the second most abundant genus. Control replicate 2, Day 17, is missing from the dataset due to the loss of bees for Day 17 sampling. *Lactobacillus mellis* and *L. mellifer* are displayed as *Bombilactobacillus mellis* and *B. mellifer* following updated classification for Firm-4 phylotype.

Principal coordinates analysis (PCoA) with Bray-Curtis dissimilarity captured 50.5% of the variation in the first two axes, PC1: 28.9%, PC2: 21.6%, for all samples. Antimicrobial treatments: Terra-Pro and tetracycline, clustered together, while ASx lysate-treated samples showed intermediate positioning between antimicrobial-treated and untreated control samples. NEWs were most distinct from all other groups, reflecting their undeveloped microbiome. Hive-reared bees formed a separate cluster from controls, treatments, and the comb, indicating environment shapes community structure (Figure 9).

**Figure 9.**
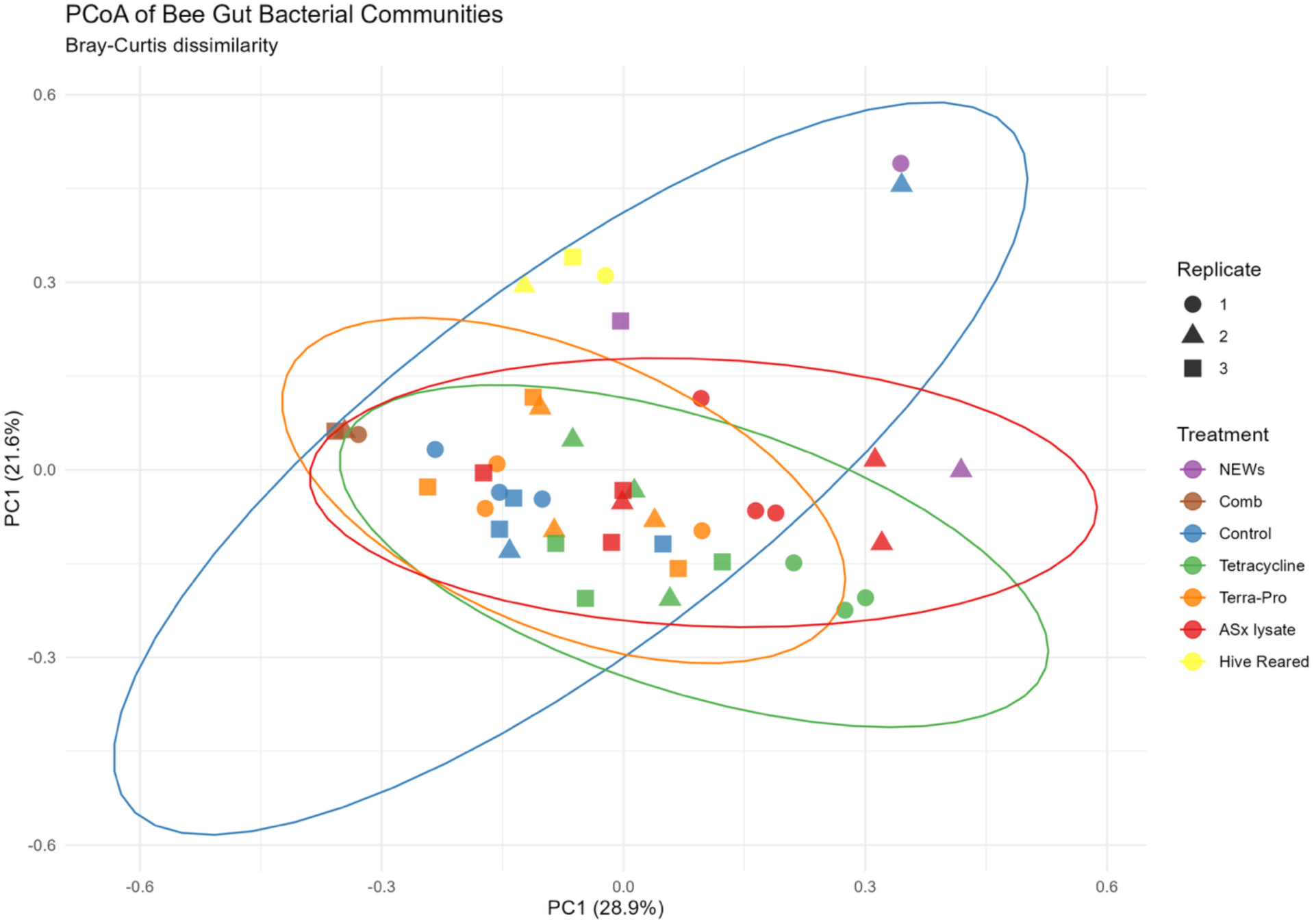
Principal coordinates analysis (PCoA) ordination of bacterial community composition using Bray-Curtis dissimilarity. Each point represents an individual sample, colored by treatment group and shaped by replicate number. Ellipses indicate 95% confidence intervals for each treatment group. PC1 and PC2 capture 28.9% and 21.6% of the variation, respectively.

Antimicrobial treatments: tetracycline and Terra-Pro cluster together, indicating similar effects on community structure. ASx lysate-treated samples show intermediate positioning between antimicrobial-treated and untreated control samples. NEWs (Newly Emerged Workers) are distinct from all other groups, reflecting undeveloped baseline microbiomes. Hive-reared bees form a separate cluster from all treatments, indicating the natural hive environment shapes community structure independent of experimental treatment effects Jaccard distance analysis revealed antimicrobial treatments were most similar, 0.27, while ASx lysate remained similar to controls, 0.37. The NEWs were most distinct from all groups, 0.65-0.72. Hive-reared bees differed from caged bees, 0.43 – 0.69, confirming rearing environment influences microbiome composition beyond initial colonization (Table-3).

DESeq2 and ANCOM-BC2 were used to identify treatment-dependent differences in bacterial taxa abundance between treatments. Compared to untreated controls, tetracycline treatment impacted 9 taxa, Terra-Pro altered 4, and ASx lysate altered only 1 taxon (DESeq2) (Supplemental file: deseq2_sig_taxa). ANCOM-BC2, a more conservative compositional method, identified no significantly different taxa at q<0.05, with one borderline taxon (*Lactobacillus kullabergensis* (q = 0.058) in the tetracycline versus control comparison (Supplemental file: deseq2_v_ ancombc2). In the DESeq2 identified taxa, tetracycline also impacted the abundance of honeybee gut-specific *Lactobacillus* species (*L. kullabergensis* and *L. apis*) (Supplemental file: deseq2_v_ancombc2).

An analysis of the comb samples placed in all the cages revealed that 50-62% of the control and treatment bee gut microbiome were also present in the comb (Supplemental file: comb_seeding). In comparison, hive-reared bees shared 91.7% of their taxa with the comb material used in caging experiments, despite exposure to natural hive comb (Supplemental file: comb_seeding).

## DISCUSSION

This study systematically demonstrated that hive-associated *Bacillus* species can improve the survival of honeybee larvae during *P. larvae* infection. Firstly, we isolated bacteria from AFB-infected combs, sequenced them, and assessed their capability to inhibit *P. larvae* growth *in vitro*. Several *Bacillus* species were identified, of which five showed inhibitory effects. Further analysis of the lysate of *B. licheniformis* revealed that its main antibacterial compound has a molecular weight between 30-50 kDa and consists of heat-stable proteins. Secondly, two *Bacillus* strains, one in culture and the other in lysate, were selected for infection trials with *P. larvae* using laboratory-reared honeybee larvae. Both enhanced larval survivability by 42-71%. Thirdly, the best *Bacillus* candidate was selected to investigate its impact on adult bee survival. We measured its impact on bee survival and microbiome composition using laboratory-based cage experiments with newly emerged adult worker bees. The survival percentage of the *Bacillus*-treated bees was significantly different from that of the control bees; however, not significantly different from that of the Terra-Pro-treated bees, a Terramycin-medicated product, used by growers to treat and control AFB pathogens. The adult gut microbiome composition of *Bacillus*-treated bees was similar to that of the controls, with minimal effects on the bee gut microbiome. We concluded that the hive-associated *Bacillus* species have the potential to improve the survival of honeybee larvae during pathogen infection, and they have minimal impacts on the adult bees, so they could be used in the control of AFB disease.

The *Bacillus* species are not core members of the honeybee gut microbiome; however, they are commonly found in the beehive, honey, pollen, and hive materials (Genersch 2010). They are reported to be detected in small numbers in bee guts (Al-Ghamdi et al., 2018). Numerous studies from different countries have shown that several *Bacillus* strains isolated from different apiarian sources exhibit antagonistic effects against various *P. larvae* strains *in vitro*. For example, *B. megaterium*, *B. licheniformis*, and *B. cereus*, isolated from honey samples from Argentina, were antagonists against *P. larvae* strains from different geographical origins (Alippi & Reynaldi, 2006; Minnaard & Alippi, 2016). *B. licheniformis* and *B. subtilis*, isolated from the guts of native honeybees in Saudi Arabia, also demonstrated *in vitro* antagonistic potentials against *P. larvae* (Al-Ghamdi et al., 2018, 2020). Different *Bacillus* species isolated from honeybee larvae showed similar antagonistic interactions *in vitro* (D Evans & Armstrong, 2005). *B. subtilis,* isolated from the honeybee gut and honey in Argentina, inhibited *P. larvae* strains (Sabaté et al., 2009). Another study from Argentina showed that *B. licheniformis* and *B. subtilis*, isolated from honey and honeybees, inhibited *P. larvae in vitro*. *B. licheniformis* showed inhibition against *P. larvae* genotypes ERIC I and IV, and *B. subtilis* inhibited *P. larvae* genotypes ERIC I, II, and IV (Bartel et al., 2019). *Bacillus* species isolated from the gut of Japanese honeybee demonstrated strong inhibitory activity against *P. larvae* (Yoshiyama & Kimura, 2009). Moreover, *Bacillus* species from sources other than typical apiarian environments, or with unreported origins, also showed inhibition against *P. larvae in vitro*. For example, *B. licheniformis* and *B. megaterium*, from GenBank in Poland, with proven probiotic potential, inhibited *P. larvae in vitro* Nowotnik et al., 2024). *B. amyloliquefaciens* LBM 5006, isolated from native soil of the Brazilian Atlantic forest, exhibited antagonistic activity against *P. larvae* (Benitez et al., 2012). *B. thuringiensis* subsp*. entomoidus*, from the USDA GenBank, showed inhibition against *P. larvae* in *in vitro* assays (Cherif et al., 2008). These results harmonize with our findings suggesting that hive-associated *Bacillus s*pecies can suppress *P. larvae* strains *in vitro*. Moreover, we also tested a *Lactobacillus* strain from the NCIMB Genbank, originally isolated from bee guts, and this strain inhibited all the tested *P. larvae* strains (Figure 3). The prior findings reported similar results: several *Lactiplantibacillus plantarum* strains (formerly *Lactobacillus plantarum*) exhibited antagonist activity against *P. larvae* ((Iorizzo et al., 2020). Likewise, two *Lactobacillus* phylotypes (Hma11 and Biut2) demonstrated strong inhibitory effects on the *in vitro* growth of *P. larvae* (Forsgren et al., 2010). Other lactic acid bacteria from different sources, such as fermented food products, can also inhibit *P. larvae* (Yoshiyama et al., 2013).

Our results of the lysate of the strain *B. licheniformis* ONT-R2-ASx against *P. larvae* suggested the presence of an antimicrobial compound with a molecular weight ranging from 30-50 kDa (Figures 3, Table 1, Figure 4). The additional tests revealed that the antimicrobial activity was mediated by its proteins, which were also heat-stable (Table 2; Figure 5). There have been no reports, to the best of our knowledge, of *B. licheniformis*-associated proteins tested against AFB pathogens or *P. larvae* strains. In fact, there are only a few reports in the literature on apiary-sourced or other *Bacillus*-associated proteins and/or bacteriocins tested against AFB pathogens. Among the apiary-sourced *Bacillus* strains, a study from Argentina reported that *B. subtilis* isolated from the honeybee gut and honey produces an antimicrobial compound, surfactin, which inhibited *P. larvae in vitro* (Sabaté et al., 2009). Similarly, bacteriocin-like inhibitory substances (BLIS) obtained from *Bacillus cereus* isolated from honey have high inhibitory effects on *P. larvae* growth, and were also heat-stable at 70 °C (Minnaard & Alippi, 2016). A 2019 study claimed that a bacteriocin called laterosporilin, a protein fraction from *Brevibacillus laterosporus* (formerly *Bacillus laterosporus*) culture supernatant, a strain isolated from the honeybee body, inhibited *P. larvae* in *in vitro* experiments (Marche et al., 2019). The non-apiarian sourced *Bacillus* strains, such as *B. amyloliquefaciens* LBM 5006, isolated from the Brazilian soil, were reported to produce an antimicrobial factor, inturin, which inhibited *P. larvae*. This inturin was reported to be heat-stable for up to 80 °C and was sensitive to proteinase K, and its exposure to the *P. larvae* cell culture resulted in a reduction in optical density associated with cell lysis (Benitez et al., 2012), which aligns with our results. Similar results were presented by Cherif et al. (2008), where a bacteriocin, entomocin 110, produced by *B. thuringiensis* subsp*. entomoidus* showed inhibitions against *P. larvae in vitro*. Like our study, entomocin 110 was heat-stable, and its inhibitory activity was completely lost after treatment with proteinase K, indicating its proteinaceous nature. However, in our study, we only used the ASx lysate with a molecular weight ranging between 30-50 kD; the specific compound or metabolites responsible for the inhibition, as well as their concentration used, were lacking. Therefore, further experiments are needed to fully characterize the specific metabolites or compounds, which could be done by mass spectroscopy and high-performance liquid chromatography analysis.

Although there are no reports of *B. licheniformis*-associated proteins tested against AFB pathogens, this strain is considered a promising probiotic for apiculture because it can enhance honeybee immunity and protect against pathogens. It has been reported that the supplementation of the normal bee diet with *B. licheniformis* and other beneficial bacteria significantly reduced the mortality of AFB-infected larvae (Al-Ghamdi et al., 2018), which has been linked to increased survival rates in honeybee larvae. Our study also showed similar results (Figure 6); however, the only difference was that Al-Ghamdi et al., (2018) study used *B. licheniformis* cell cultures, whereas we used the lysate (50-30 kDa) to feed the larvae. It may be possible that antimicrobial peptides (e.g., bacteriocins) associated with this strain may have played a role in the mortality reduction of the AFB-infected larvae in the study conducted by Al-Ghamdi et al., (2018). However, this is just a presumption; to uncover the real underlying mechanisms leading to the mortality reduction in the infected larvae in such infection studies, more investigations are necessary, such as scanning electron microscopy, confocal microscopy in combination with Mass spectroscopy and high-performance liquid chromatography, as mentioned earlier.

Apart from Al-Ghamdi et al. (2018), no other reports mention *B. licheniformis* against AFB diseases in honeybees. Nevertheless, *B. licheniformis* has been reported to have significant probiotic potential in animal and human health in terms of modulation of the intestinal microbiota, antimicrobial activity, growth promotion, anti-inflammatory and immunostimulatory effects, promotion of the regulation of the lipid profile, increase of neurotransmitters, and stress reduction (Ramirez-Olea et al., 2022). A study by Díaz et al.(2022) stated that lipopeptides synthesized by the *B. licheniformis* B6 strain have potential antibacterial and anti-biofilm activity against pathogenic bacteria of health importance, suggesting it as a potential biocontrol alternative. The same review paper Ramirez-Olea et al.(2022) also mentioned that 70% of the reviewed articles reported *B. licheniformis* as a probiotic, out of which 75% used it in animal trials (such as on birds, including chickens, marine animals, and rodents) or *in vivo* models. Additionally, *B. licheniformis* has proven to be a safe probiotic for consumption, with the ability to resist the conditions of the entire gastrointestinal system (Ramirez-Olea et al., 2022). Since *B. licheniformis* has proved itself safe and has already been used in the veterinary industry, it could also be used in apiculture against AFB diseases, as it has shown positive results against *P. larvae* in honeybee larvae.

Regarding the antimicrobial compound size 30-50 kDa produced by our *B. licheniformis* strain, a review paper reported that *B. licheniformis* can produce various bacteriocins ranging in molecular weight from 1.4 to 55 kDa (Shleeva et al., 2023). A 30.7 kDa, YbdN protein extracted from *B. licheniformis* showed antibacterial activity against methicillin-resistant *Staphylococcus aureus*, vancomycin-resistant Enterococci, and *Listeria monocytogenes* (Jamal et al., 2006). Likewise, antimicrobial compounds named F4, F5, and F6 with molecular mass less than 45 kDa, produced by *B. licheniformis* BFP011, showed resistance to pronase enzyme and high temperature at 100 °C and 121 °C for 15 minutes and were able to inhibit the growth of gram-negative bacteria *Escherichia coli* and *Salmonella typhi* (Arbsuwan et al., 2014). Since *B. licheniformis* possesses remarkable plasticity as a secretion vehicle, the compounds it secretes depend on the growth medium composition, growth period, environmental conditions, and the specific strain (Shleeva et al., 2023).

Apart from *B. licheniformis*, we also tested cell cultures of *B. zhangzhouensis* strain 2D and an NCIMB *Lactobacillus* strain in the laboratory-reared larvae against *P. larvae*. Our results showed that cell cultures of the *B. zhangzhouensis* strain 2D, supplemented with the larval diet, significantly reduced larval mortality by 42% (Figure 6A-F, I.2), indicating the potential of this strain against AFB infections. To the best of our knowledge, *B. zhangzhouensis* was first proposed as a new *Bacillus* strain in 2016, isolated from aquaculture water from Zhangzhou city in China (Liu et al., 2016). Later, it was reported to be isolated from honey samples (Brudzynski, 2021; Pajor et al., 2018; Pomastowski et al., 2019), bee pollen and bread (Pełka et al., 2021), from seaweeds (Shah et al., 2024), and recently from plants (Toshmatov et al., 2025). Based on our current information, this is the very first report demonstrating that *B. zhangzhouensis* has the potential to inhibit *P. larvae*, causing AFB disease in honeybees. However, there are studies that reported that *B. zhangzhouensis* produces metabolites with growth inhibition potential against human and animal pathogens, including *Staphylococcus aureus, S. epidermidis, Candida albicans, Listeria monocytogenes* (Pajor et al., 2018), and *Escherichia coli, Pseudomonas aeruginosa* (Pełka et al., 2021). A paper also reported that a phthalate derivative extracted from an epibiotic bacterium, *Bacillus zhangzhouensis* SK4, isolated from seaweeds, showed significant suppression of QS-regulated virulence factors of *P. aeruginosa* MCC 3457, which could deal with the multidrug resistance phenomenon (Shah et al., 2024). Recently, it has also been reported that *B. zhangzhouensis*, isolated from plants, produces secondary metabolites with antifungal and growth-regulating properties in plants (Toshmatov et al., 2025).

The NCIMB *Lactobacillus* strain fed alone with the larval diet showed positive results in the larval survival, suggesting that the *Lactobacillus* strain has no negative impacts on larval growth; however, the larval survival significantly decreased when it was co-inoculated in the diet with *P. larvae*. Our findings did not match our own *in vitro* results, and are contrary to the reports that demonstrated that the addition of different *Lactobacillus* species to the larval food improved honeybee larval survival during *P. larvae* infection ( Daisley et al., 2020; Forsgren et al., 2010). Though these discrepancies might be explained by the fact that Forsgren et al. (2010) and Daisley et al. (2020) administered a mixture of lactic acid bacteria (LAB), including different *Lactobacillus* species, in the exposure bioassays, resulting in a significant reduction in the number of AFB-infected larvae. However, we administered only a single *Lactobacillus* strain, likely taking more competitive advantages by the *P. larvae* strain in the supplemented diets, resulting in a reduction of *Lactobacillus* cell viability. Additionally, in the study by Daisley et al. (2020), *Lactobacilli* strains were delivered through a nutrient biopatty instead of the liquid larval feed. This might be an additional reason why the supplemented *Lactobacillus* strain, in combination with the pathogenic strain in the diet, failed to demonstrate growth suppression of *P. larvae*, contradicting the *in vitro* assay results. We did observe a trend towards increased survival, but the increase was not significant (P=0.1).

Honeybees are characterized by a uniquely conserved gut microbial community consisting of 5-8 bacterial phyla that support diverse host functions from metabolic regulations, cognitive enhancement, nutrient processing, to priming the host immune system (Engel et al., 2015; Han et al., 2024; Zheng et al., 2019). Gut symbionts colonize the hindgut in a successive order within the first 4-8 days post NEW eclosion. If the microbial community abundance fluctuates across the insect’s lifespan, disruptions such as species loss or dysbiosis can result (Motta & Moran, 2024). Therefore, any effective treatment strategies against bacterial pathogens must account for impacts on both microbiome development and the established adult microbial community. Previous studies on the honeybee gut microbiome establishment reported that NEWs acquire their characteristic gut bacteria through exposure to hive components, horizontal transmission from hive mates, and coprophagia during early adult development (Martinson et al., 2012; Powell et al., 2014). As traditional *in vivo* hoarding cages lack these natural transmission routes, brood comb from the hive was provided to each cage to facilitate microbiome establishment in the absence of the hive environment.

To assess the impact of treatment and early exposure to ASx lysate on the bee gut microbiome, gut microbial communities were analyzed using 16S rRNA gene sequencing and compared to those treated with commonly administered antimicrobials, Terra-Pro and tetracycline. We found that the survival of *Bacillus*-treated bees was slightly lower than that of the sugar-syrup-fed bees; however, no survival difference between the Terra-Pro-treated bees, the antibiotics that the growers use to treat AFB in field conditions. Beta diversity analysis of the relative abundance using Bray-Curtis dissimilarity showed ASx lysate treatment had minimal impact on microbiome composition when compared to untreated controls and produced similar alterations to tetracycline and Terra-Pro (Figure 9). DESeq2 differential abundance analysis identified treatment-specific changes for all treatments; however, only tetracycline induced a change in *Apis*-specific *Lactobacillus* species abundance by enriching *L. kullabergensis* and *L. apis* while decreasing *L. bombicola* (Supplemental file: deseq2_sig_taxa). ANCOM-BC2, a more conservative compositional method, identified zero significant taxa at q < 0.05, with one borderline result in the tetracycline treatment for *L. kullabergensis* (Supplemental file: ancombc2 borderline). The enrichment of specialist species following antibiotic treatment may reflect competitive release following suppression of other community members as opposed to stimulation. Critically, ASx lysate showed no significant abundance changes in core microbiome bacterial species in either analytical method, supporting its potential as a treatment option for AFB disease control. These results suggested that ASx lysate may offer a targeted approach that minimizes disruption to the beneficial core microbiome.

## CONCLUSION

In this study, we presented a systematic investigation to suggest that the honeybee hive harbors bacteria, specifically *Bacillus* species, that improve the survival of honeybee larvae during pathogen infection. Specifically, we isolated hive-associated bacteria, taxonomically identified them using 16S sequencing, and showed that several *Bacillus* species, either their bacterial cultures, lysates, or both, have the potential to inhibit one or more *P. larvae* strains, both *in vitro* and in larval infection trials. For the infection trials, we used laboratory-reared honeybee larvae because larvae are susceptible to *P. larvae* pathogens, causing AFB disease. The lysate of *B. licheniformis* has an antimicrobial compound with a molecular weight between 30-50 kDa, with heat-stable proteins, responsible for the inhibitory effects against all tested *P. larvae* strains.

Finally, we conducted adult cage experiment with newly emerged bees to determine whether the lysate of one hive-associated bacterium, *B. licheniformis*, affects the survival rate and the bee gut microbiome. The survival rate of lysate-treated bees was not significantly different than the Terra-Pro-treated ones, and the microbiome composition of the lysate-treated bees was similar to the control treatments and had minimal effects on the bee gut microbiome. These results infer that the hive-associated *Bacillus* species could improve the honeybee larval survival during pathogen infection and could be used as an antimicrobial therapy to control AFB disease.

## ACKNOWLDEGEMENT

We thank Zoe Wilson and Lucelia De Moura Pereira (University of Idaho) for their support with apiary work and the cage experiment. We thank Dr. Christine Parent and her lab members (University of Idaho) for providing their lab space for conducting our cage experiment. We thank Dr. Paul A. Rowley (University of Idaho) for helpful suggestions about the bacterial lysate work. We thank Dr. Brandon Hopkins (Washington State University) for providing us with AFB-infected comb samples. The research reported in this publication was supported by the National Institute of General Medical Sciences of the National Institutes of Health under Award Number P20GM103408 and the National Institute of Food and Agriculture of the US Department of Agriculture under Award Number 2023-67013-39067.

## Supplementary Figure legend

**Figure S1.**Representative plates of spot plate assay of heat-treated ONT-R2-ASx (50-30 kDa) lysate on the lawns of *P. larvae* strains. The lysate was serially diluted (10^0^ to 10^-3^), heat-treated, and spotted onto the lawns.

